# Anion exchange regulates intestinal mucus expansion while Cftr regulates *de novo* mucus secretion

**DOI:** 10.1101/2024.03.18.585614

**Authors:** Penny L Ljungholm, Anna Ermund, Molly M Söderlund Garsveden, Victor L Pettersson, Jenny K Gustafsson

**Author notes:** Correspondence should be addressed to: Jenny Gustafsson, University of Gothenburg, Department of Physiology, Medicinaregatan 11, Box 432, 405 30 Gothenburg, Sweden.

## Abstract

The intestinal epithelium is covered by mucus that protects the tissue from the luminal content. Studies have shown that anion secretion via the cystic fibrosis conductance regulator (Cftr) regulates mucus formation in the small intestine. However, mechanisms regulating mucus formation in the colon are less understood. The aim of this study was to explore the role of anion transport in regulation of mucus formation during steady state, and in response to carbamylcholine (CCh) and prostaglandin E_2_ (PGE_2_). CftrΔF508 (CF) mice were used to assess the role of Cftr, and 4,4′-diisothiocyanatostilbene-2,2′-disulfonate (DIDS) was used to inhibit anion exchange. In the distal colon, steady state mucus expansion was reduced by apical DIDS, and normal in CF mice. PGE_2_ stimulated mucus expansion without *de novo* mucus secretion in wild type (WT) and CF distal colon via DIDS sensitive mechanisms, while CCh induced *de novo* mucus secretion in WT but not in CF colon. However, when added simultaneously, CCh and PGE_2_, stimulated *de novo* mucus secretion in CF colon via DIDS sensitive pathways. A similar response was observed in CF ileum that responded to CCh and PGE_2_ with DIDS sensitive *de novo* mucus secretion. In conclusion, this study suggests that apical anion exchange regulates intestinal mucus expansion, while Cftr regulates *de novo* mucus secretion from ileal and distal colon crypts. Furthermore, these findings demonstrate that in the absence of a functional Cftr channel, activation of anion exchange can help release mucus from intestinal goblet cells.

## Introduction

The intestinal epithelium is a dynamic structure capable of alternating between absorption and secretion to fine tune ion and water homeostasis, and to maintain a protective barrier against the large amount of microorganisms that reside in our gastrointestinal tract [1-3]. Coordinated secretion of ions, fluid, and mucus each contribute to maintaining the intestinal barrier. Anion secretion provides the driving force for fluid secretion, and regulates the ionic milieu required for normal mucus layer formation [4-6]. Building of the intestinal mucus barrier is a multi-step process starting with exocytosis of the mucin granules, followed by expansion and processing of the secreted mucins to form the mucus barrier [7-9]. In the small intestine, the mucus forms a loose and permeable structure that acts as a diffusion barrier for antimicrobial proteins and peptides and a habitat for the microbiota [10-13]. The loose structure of the small intestinal mucus makes it easily transported in the distal direction which combined with the secretion of antimicrobial proteins and peptides keep the amounts of bacteria lower in the small intestine as compared to the colon [14]. In the mid and distal colon of mice, the mucus forms a dense adherent barrier that separates the vast majority of the commensal microbiota from accessing the underlying epithelium [1, 15, 16]. Thus, the properties of the mucus layers differs between the small intestine and colon despite the fact that the main structural component of the mucus layer, mucin 2 (Muc2), is shared by the two organs [17]. The physiological importance of secreting mucus with the correct properties is exemplified in the disease cystic fibrosis where loss of a functional cystic fibrosis conductance regulator (CFTR) that transports chloride and bicarbonate results in mucus plugging in CFTR expressing organs such as the airways, small intestine, pancreas and reproductive tract, which in turn leads to inflammation, and subsequent organ damage [18, 19]. Studies in the small intestine and airways of mice and pigs have shown that loss of Cftr mediated anion transport, in particular bicarbonate, renders the small intestinal and airway mucus into a dense and adherent layer that acts as a binding point, and nutrient source for bacteria, resulting in bacterial overgrowth [4, 9, 20, 21]. In the colon, where the mucus normally is dense and adherent, loss of CFTR mediated transport causes less severe pathology as compared to the small intestine, indicating that additional transport processes are involved in regulating mucus layer formation in the colon [22]. Studies have shown that Tmem16A a Ca^2+^ regulated anion channel, is involved in regulating baseline mucus release in the mouse colon, however whether other anion transporters such as anion exchangers regulate other aspects of mucus layer formation such as expansion of the mucus following exocytosis, or *de novo* mucus secretion in response to secretagogues is not known [23]. In the present study we explored the role of the Cftr and anion exchange in regulation of mucus layer formation during steady state and in response to the muscarinic receptor agonist carbamylcholine (CCh) and prostaglandin E_2_ (PGE_2_). To assess the role of the Cftr and anion exchange in regulation of mucus layer formation we measured mucus growth over time and mucus adhesion in ileal and colonic intestinal explants *ex vivo*. We have previously shown that in distal colon but not in ileum explants, mucus grows at a steady state rate of ∼2 µm/min, a process involving both expansion of previously secreted mucus and *de novo* mucus secretion [24]. To be able to distinguish between mucus expansion and *de novo* mucus secretion in the distal colon, we first measured mucus growth over time, and then processed the tissue for immuno-histological analysis to evaluate the mucus content of the tissue.

## Materials and Methods

### Mouse tissue

The experiments were performed using male and female homozygous Cftr F508 (CF) and wild type littermate controls on a C57/BL6 background [25]. Mice were housed in a specific pathogen-free facility, fed a routine chow diet, had access to water and food *ad libitum*, and kept at a 12h light/dark cycle at a constant temperature (21-22°C). Animals were between 8 and 15 weeks old at the time of the experiments. To increase the survival of the homozygous Cftr F508 mice, they were given an osmotic laxative in the drinking water (PEG4000 18 mM, KCl 10 mM, Na_2_SO_4_ 40 mM, NaHCO_3_ 84 mM and NaCl 25 mM). The animals were transferred to normal tap water 3 days prior to the experiments.

### Mucus thickness measurements

Measurements of mucus thickness were performed as described previously using apical Kreb’s mannitol and basal Kreb’s glucose buffer [26]. Briefly, the distal colon or ileum was dissected, flushed with ice-cold oxygenated Krebs’ buffer and kept on ice for 30 min. Following incubation, the tissue was flushed once more with Krebs’ buffer, opened along the mesenteric border and the longitudinal muscle layer was removed by blunt dissection. The distal colon or ileum was divided into two parts and studied in parallel. To visualize the mucus layer a suspension of activated charcoal particles was added to the apical surface and allowed to sediment down to the top of the mucus layer. The thickness of the distal colon mucus layer was assessed by measuring the distance between the mucus surface and the epithelial surface using a micropipette connected to a micromanipulator and a digimatic indicator. In the ileum experiments, the tissue was mounted in the chamber, followed by removal of the mucus by aspiration or scraping. The thickness of the remaining mucus was then measured by measuring the distance between the crypt openings and the mucus surface. Thus, in the ileum the mucus thickness at t0, represents the mucus that is left in between the villi following removal. Carbachol (CCh) (1 mM), Prostaglandin E_2_ (PGE_2_) (10 µM), and the combination of CCh and PGE_2_ were added to the basal buffer, and the effect on mucus thickness was evaluated after 15/30 min in the colon, and 20/40 min in the ileum. In the inhibitory experiments, DIDS (apical or basal) and the apical chloride free buffer were added 30 min prior to CCh and PGE_2_ stimulation in the colon, and 20 min prior to CCh and PGE_2_ stimulation in the ileum.

### Lectin staining

Distal colon tissues from the mucus measurement experiments were fixed in the measurement chamber for 1h using 4 % buffered formaldehyde (Histolab, Askim, Sweden), transferred to a tube containing fresh fixative and incubated over-night, followed by paraffin embedding and sectioning. 4 µm thick sections were deparaffinized, rehydrated, and stained with fluorescein conjugated UEA1 (25 µg/ml) (cat. no: FL-1061, Vector Laboratories, Burlingame, CA) and AlexaFluor 647 conjugated WGA (5 µg/ml) (cat no: W32466, Thermo Fisher Scientific, Waltham, MA) for 30 min followed, by 5 min wash in PBS and counterstained with Hoechst DNA stain for 5 min and imaged using a Nikon microscope. Acquired images were analyzed using Fiji [27] and Imaris (Bitplane, Belfast, Great Britain) software. A combination of UEA1 and WGA was used to stain the entire goblet cell population as UEA1 stains the surface goblet cells and goblet cells in the upper half of the crypt, while WGA stains goblet cells in the lower half of the crypt.

### Quantification of mucus secretion

Mucus secretion in the surface and crypt epithelium was quantified using Fiji [27] by measuring the sum of the theca areas of surface or crypt goblet cells, divided by the area of either the surface or crypt area. To quantify the mucus content of the surface epithelium, the sum of the theca areas of all UEA1^+^ inter-crypt goblet cells was measured and divided by area of the surface epithelium. To quantify the mucus content of the colonic crypts, sum of the theca areas of all UEA1^+^ and/or WGA^+^ crypt goblet cells was divided by the area of the crypt cross section. Both the surface and crypt mucus content are expressed as percentage. Mucus secretion was defined as a significant decrease in mucus content of the respective compartment.

### Drugs and buffer composition

Carbamylcholine chloride (CCh) (cat no: C4382, P0409, Sigma-Aldrich, Steinheim, Germany) was dissolved in water. Prostaglandin E_2_ (PGE_2_) was dissolved in 50:50 ethanol and DMSO. 4,4′-diisothiocyanatostilbene-2,2′-disulfonate (DIDS) (cat no: S347523, Sigma-Aldrich, Steinheim, Germany) was dissolved in DMSO. The Krebs’ buffer had the following composition in mM: NaCl 115.8, CaCl_2_ 1.3, KCl 3.6, KH_2_PO_4_ 1.4, NaHCO_3_ 23.1, and MgSO_4_ 1.2 (Merck, Darmstadt, Germany). The Krebs-mannitol buffer also contained Na-Pyruvate (5.7 mM) (Sigma-Aldrich, Steinheim, Germany), Na-L-Glutamate (5.1 mM) (Merck, Darmstadt, Germany) and D-Mannitol (10 mM) (Sigma-Aldrich, Steinheim, Germany), and the Kreb-glucose buffer contained Na-Pyruvate (5.7 mM), Na-L-Glutamate (5.1 mM) and D-Glucose (10 mM) (Sigma-Aldrich, Steinheim, Germany). In the chloride free experiments NaCl, KCl and CaCl_2_ were replaced with equimolar concentrations of Na-gluconate and K-gluconate, and twice the concentration of Ca-gluconate (Sigma-Aldrich, Steinheim, Germany) to compensate for the chelation of calcium by gluconate. pH was set to 7.4 using acetic acid.

### Statistics

Data are presented as mean ± standard error of the mean (SEM). Single comparisons between two groups were made using Mann Whitneys’ test. Comparisons between two groups over time were made using a two-way ANOVA with Sidaks’ post-hoc test. Comparisons between three groups or more were made using Kruskal-Wallis test with Dunns’ post-hoc test. A p-value <0.05 was considered statistically significant.

## Results

### Steady state mucus expansion in the distal colon is mediated by apical anion exchange

Previous studies have shown that in the small intestine, loss of Cftr mediated bicarbonate secretion renders the normally loose and permeable mucus into a dense adherent structure [8]. To explore the role of Cftr in regulation of mucus formation in the colon, we measured baseline mucus thickness and growth over time in the distal colon of C57/BL6 (WT) and Cftr 508 (CF) mice, and evaluated the mucus content of the tissues by lectin staining. Our results showed no difference in either the initial mucus thickness or baseline mucus growth comparing WT and CF distal colon (Fig. 1A). Analysis of the mucus content of the tissue showed no difference between WT and CF mice in either the surface epithelium (Fig. 1B, representative pictures in C) or crypt epithelium (Fig. 1C, representative pictures in C).

**Fig. 1:**
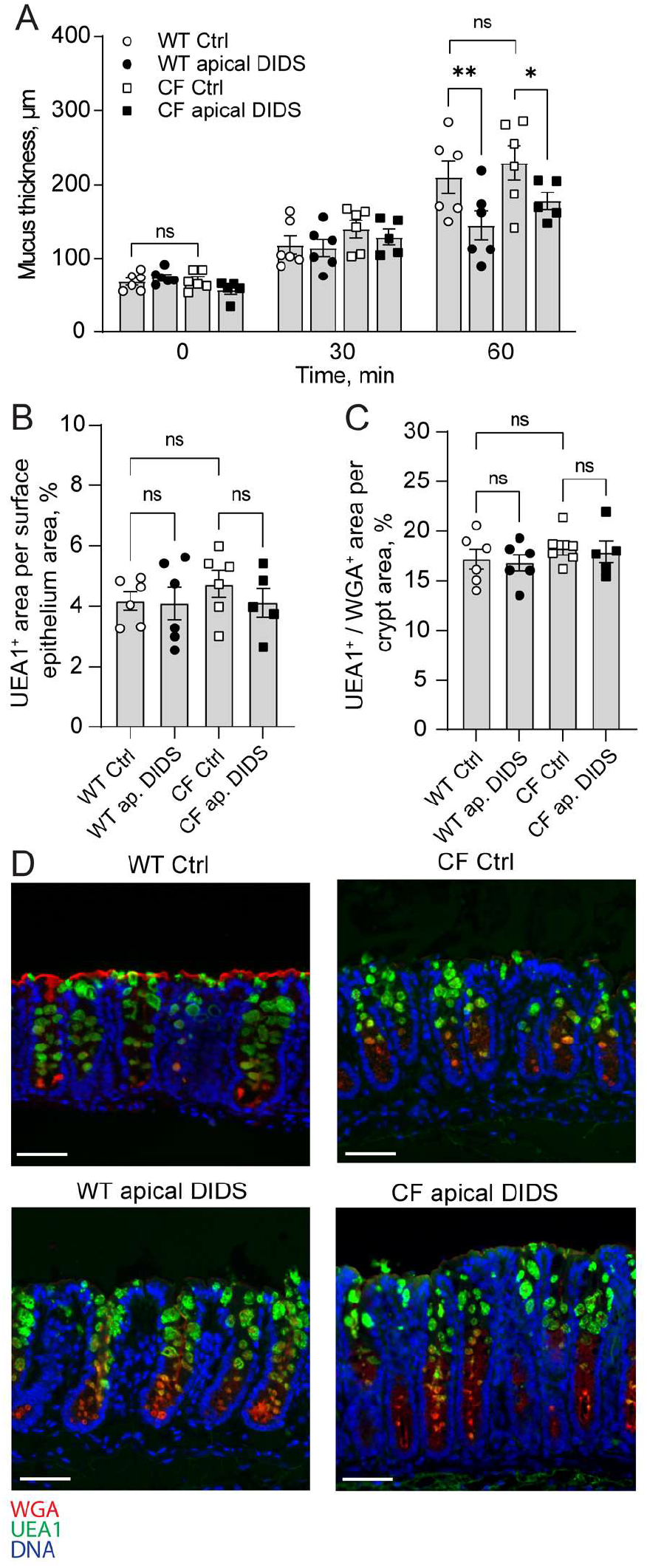
Steady state mucus expansion in the distal colon is mediated by apical anion exchange. A) Mucus thickness over time in WT ctrl (n=6), WT treated with apical DIDS (200 µM) (n=6) at t0, CF ctrl (n=6) and CF treated with apical DIDS at t0 (n=5). B) UEA1^+^ mucus area per epithelial surface area. C) UEA1^+^/WGA^+^ mucus area per distal colon crypt area. D) Representative wide-field fluorescent images of WT and CF distal colon. Scale bar: 100 µM. Data are presented as mean ± SEM, ^*^p < 0.05, ^**^p < 0.01, n.s = non-significant. Statistical analysis was performed using a two-way ANOVA with Sidaks post-hoc test (A) and Kruskal-Wallis test with Dunn’s post-hoc test (B).

In addition to the Cftr, the distal colon expresses anion exchangers that take up Cl^-^ and release bicarbonate into the colonic lumen [28]. To explore whether apical anion exchange is involved in regulating baseline mucus formation in the distal colon, we repeated the experiments in the presence of the anion-exchange inhibitor 4,4′-diisothiocyanatostilbene-2,2′-disulfonate (DIDS). The results showed that DIDS (200 µM) added to the apical buffer reduced baseline mucus growth in both WT and CF distal colon (Fig. 1A). Analysis of the mucus content of the tissue showed no effect of DIDS in either the surface or crypt epithelium, respectively (Fig. 1B, representative pictures in D). Together these results demonstrate that steady state mucus formation in the distal colon is independent of a functional Cftr channel but dependent on DIDS sensitive apical anion exchange.

### Effect of PGE_2_ and CCh in regulation of mucus layer formation in WT and CF distal colon

To further characterize the role of anion transport in regulation of mucus formation in the distal colon we exposed WT and CF tissues to two secretagogues, the acetylcholine agonist CCh and PGE_2_. We assessed the effect of the two substances on mucus layer formation by measuring changes in mucus growth rate, and mucus content of the tissue. In WT mouse distal colon, exposure to CCh triggered a transient increase in mucus growth rate, induced a reduction in the mucus content of the crypt epithelium, and reduced the number of mucus containing crypt goblet cells, indicative of complete emptying of a subset of goblet cells (Fig. 2A-C, representative pictures in D). Stimulation of WT distal colon tissues with PGE_2_ induced a transient increase in mucus growth rate, similar to that observed in response to CCh (Fig. 2A). However, PGE_2_ did not affect the mucus content of the tissue, or the number of mucus containing goblet cells (Fig. 2B-C, representative picture in D), suggesting that PGE_2_ regulates mucus expansion rather than *de novo* mucus secretion.

**Fig. 2:**
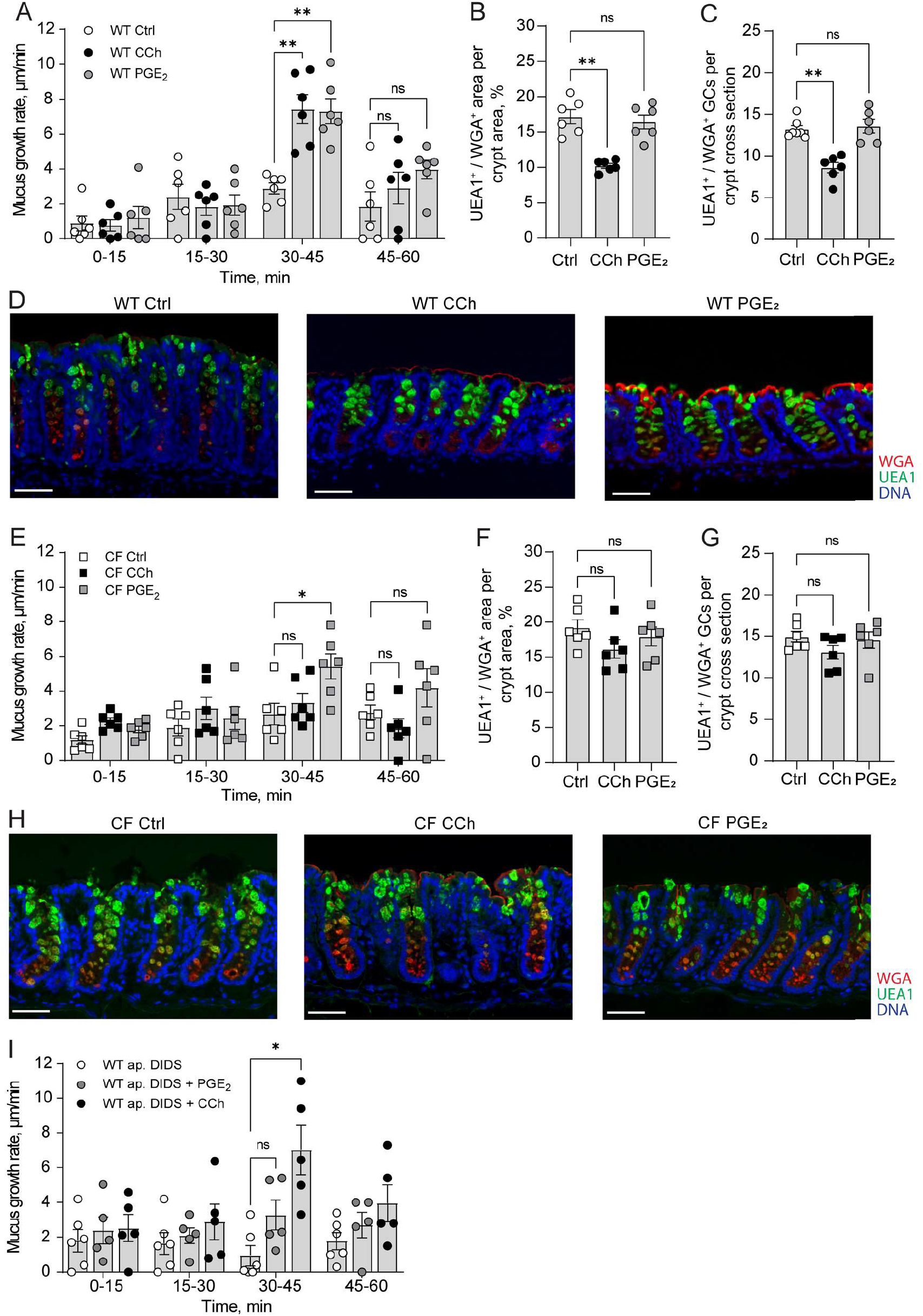
Effect of PGE_2_ and CCh in regulation of mucus layer formation in WT and CF distal colon. A) Mucus growth rate in WT ctrl (n=6), WT treated with CCh (1 mM) (n=6) at t30, WT treated with PGE_2_ (10 µM) (n=6) at t30. B) UEA1^+^/WGA^+^ mucus area per distal colon crypt area. C) Number of UEA1^+^/WGA^+^ goblet cells per distal colon crypt cross section. D) Representative wide-field fluorescent images of WT distal colon. E) Mucus growth rate in CF ctrl (n=6), CF treated with CCh (1 mM) (n=6) at t30, CF treated with PGE_2_ (10 µM) (n=6) at t30. F) UEA1^+^/WGA^+^ distal colon mucus area per crypt area. G) Number of UEA1^+^/WGA^+^ goblet cells per distal colon crypt cross section. H) Representative wide-field fluorescent images of CF distal colon. I) Mucus growth rate in WT distal colon treated with apical DIDS at t0 (200 µM) (n=6), WT distal colon treated with apical DIDS at t0 and PGE_2_ (10 µM) at t30 (n=5), and WT distal colon treated with apical DIDS at t0 and CCh (1 mM) at t30 (n=5). Scale bar: 100 µM. Data are presented as mean ± SEM, ^*^p < 0.05, ^**^p < 0.01, n.s = non-significant. Statistical analysis was performed using a two-way ANOVA with Sidaks’ post-hoc test (A, E and I) and Kruskal-Wallis test with Dunn’s post-hoc test (B, C, F and G).

In CF mouse distal colon, exposure to CCh failed to induce an increase in mucus growth rate (Figure 2E), and no effect was observed with respect to the mucus content of the tissue, or the number of mucus containing goblet cells (Figure 2F-G, representative pictures in H). In contrast, PGE_2_ induced a transient increase in the mucus growth rate, similar to that observed in WT distal colon (Fig. 2E). PGE_2_ did not affect the mucus content of the tissue or the number of mucus containing goblet cells in CF distal colon, similar to that observed in WT distal colon (Fig. 2F-G, representative pictures in H).

Based on our results demonstrating that baseline mucus growth was regulated by DIDS sensitive apical ion transport, we proceeded to explore whether the PGE_2_ induced effect on mucus growth was regulated by similar mechanisms. Our results showed that pretreatment of WT distal colon with apical DIDS inhibited the PGE_2_ effect on mucus growth rate but had no effect on the CCh induced increase in mucus growth rate (Fig. 2I). Combined these results suggest that in the distal colon, CCh induces *de novo* mucus secretion via Cftr dependent processes, while PGE_2_ induces mucus expansion via activation of DIDS sensitive apical anion transport.

### Stimulation of CF distal colon with CCh and PGE_2_ induces de novo mucus secretion

Based on our finding of an impaired mucus secretory response to CCh but an intact mucus growth response to PGE_2_ in CF distal colon, we explored whether activation of PGE_2_ mediated transport pathways could normalize the impaired CCh response. Distal colon tissues from CF mice were exposed to a combination of CCh (1 mM) and PGE_2_ (10 µM), and the effect on mucus growth rate, mucus content in the tissue, and the number of mucus containing goblet cells were measured. Stimulation of tissues with CCh and PGE_2_ induced a transient increase in mucus growth rate (Fig. 3A) that was paralleled by a decrease in the mucus content of the tissue, and a decrease in the number of mucus containing goblet cells (Fig. 3B-C, representative pictures in D). To determine whether the mucus secretory response involved activation of DIDS sensitive apical transporters, the experiments were repeated in the presence of apical DIDS. Pretreatment with apical DIDS for 30 min inhibited the CCh + PGE_2_ induced increase in mucus growth rate, inhibited the reduction in mucus content in the tissue and blocked the reduction of the number of mucus containing goblet cells (Fig. 3A-C, representative pictures in D). In contrast, 30 min pretreatment with DIDS added to the basal side of the tissue had no effect on either of the three parameters (Fig. 3A-C, representative pictures in D). To further test the type of anion exchanger involved in regulating this process we repeated the experiments using chloride free apical buffer to inhibit Cl^-^/HCO_3_^-^ exchange. Removal of chloride from the apical buffer reduced the CCh + PGE_2_ response by 90 % (Fig. 3E), pointing towards a role for Cl^-^/HCO_3_^-^ exchange in regulating this process. Combined these results suggest that activation of PGE_2_ induced Cl^-^/HCO_3_^-^ exchange can help release mucus from the goblet cells in CF distal colon.

**Fig. 3:**
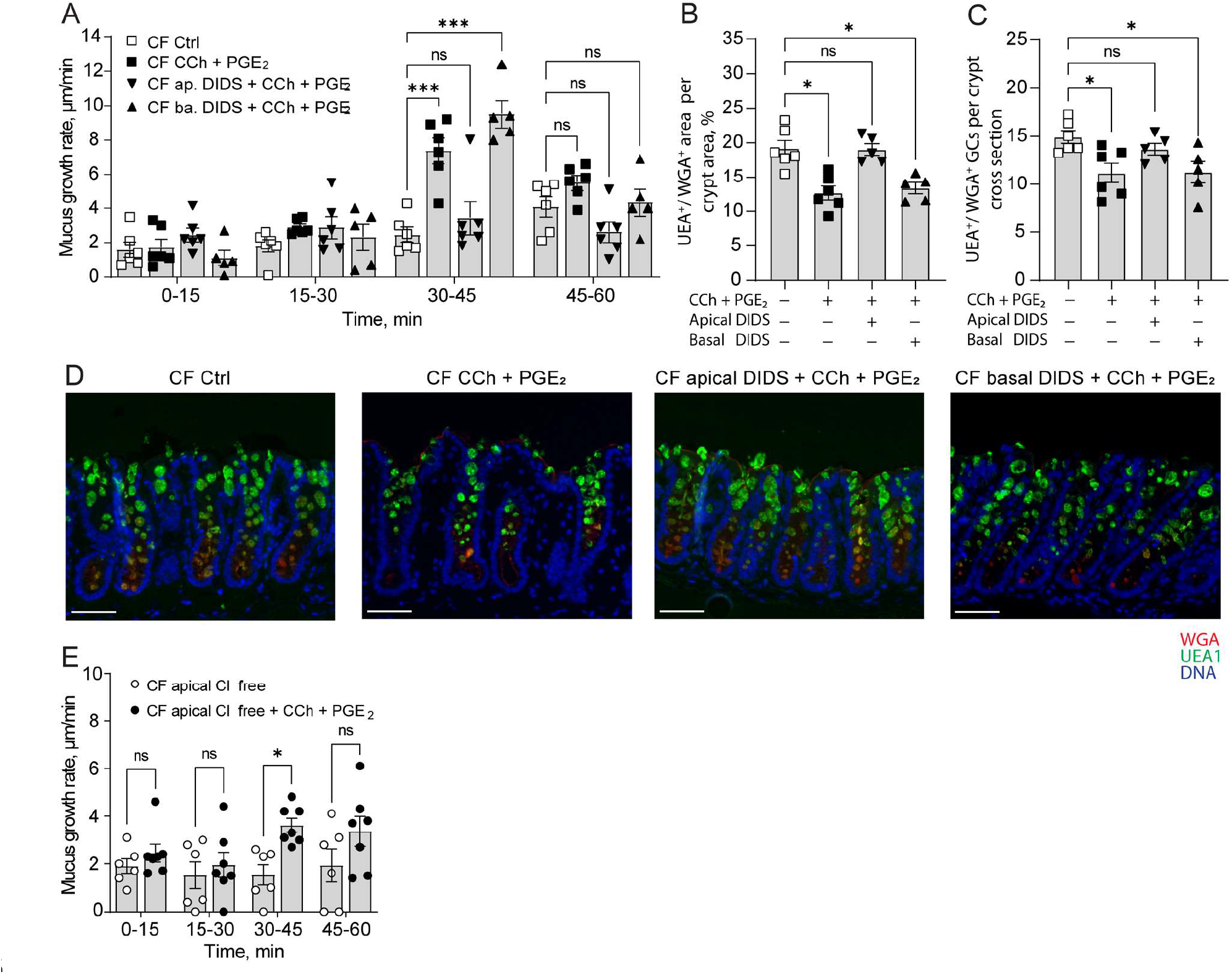
Parallel stimulation of CF distal colon with CCh and PGE_2_ induces de novo mucus secretion. A) Mucus growth rate in CF ctrl (n=6), CF treated with CCh (1 mM) and PGE_2_ (10 µM) at t30 (n=6), CF treated with apical DIDS (200 µM) at t0 followed by CCh (1 mM) and PGE_2_ (10 µM) at t30 (n=6), CF treated with basal DIDS (200 µM) at t0 followed by Ch (1 mM) and PGE_2_ (10 µM) at t30 (n=5). B) UEA1^+^/WGA^+^ distal colon mucus area per crypt cross section. C) Number of UEA1^+^/WGA^+^ goblet cells per distal colon crypt cross section. D) Representative wide-field fluorescent images of CF distal colon. E) Mucus growth rate in CF treated with apical Cl^-^ free buffer at t0 (n=6), and CF treated with apical Cl^-^ free buffer at t0 and CCh (1 mM) and PGE_2_ (10 µM) at t30 (n=7). Scale bar: 100 µM. Data are presented as mean ± SEM, ^*^p < 0.05, ^**^p < 0.01, ^***^p < 0.001, n.s = non-significant. Statistical analysis was performed using a two-way ANOVA (A and E) with Sidaks’ post-hoc test, and Kruskal-Wallis test with Dunn’s post-hoc test (B-C).

### Effect of CCh and PGE_2_ in regulating mucus layer formation in the WT and CF ileum

In the small intestine, loss of Cftr mediated transport results in mucus adhering to the tissue [8]. We have previously shown that stimulation of ileal tissues with the combination of CCh and PGE_2_ induces *de novo* mucus secretion in both WT and CF tissues, with the main differences between the groups being that the secreted mucus is more dense and adherent to the tissue [8]. Based on our findings from the distal colon, we asked whether CCh induced *de novo* mucus secretion is impaired also in the CF ileum, and whether the previously observed intact mucus secretory response to CCh and PGE_2_ relies on activation of apical anion exchange. To address these questions, we exposed WT and CF ileal explants to CCh (10 µM), PGE_2_ (10 µM) and the combination of CCh (10µM) and PGE_2_ (10µM) and measured the effect on mucus thickness over time and mucus adhesion. In the WT ileum, stimulation with CCh induced an increase in mucus thickness, while PGE_2_ had no effect on the mucus thickness (Fig. 4A). Stimulation of WT ileal tissues with the combination of CCh and PGE_2_ induced an increase in mucus thickness similar in magnitude to that observed in response to CCh (Fig. 4A). In the CF ileum, neither CCh, nor PGE_2_ induced an increase in mucus thickness when added separately (Fig. 4B). However, stimulation of CF ileum with the combination of CCh and PGE_2_ induced a significant increase in mucus thickness, similar in magnitude to that observed in WT ileum (Fig. 4B). To test whether the CCh and PGE_2_ induced increase in mucus thickness in CF ileum was mediated by anion exchange, we repeated the experiments in the presence of apical DIDS. Pretreatment with apical DIDS reduced the CCh and PGE_2_ induced increase in mucus growth in CF ileum by 84.2 % (Fig. 4B). To test the adhesive properties of the secreted mucus we aspirated the mucus and measured the remaining thickness. In the WT ileum, 57 ± 5.0 % of the CCh induced mucus, and 78 ± 3.8 % of the CCh + PGE_2_ induced mucus could be aspirated, while in the CF ileum all of the CCh + PGE_2_ induced mucus remained remained attached to the tissue (Fig. 4C). Combined these results suggest that in the ileum, CCh induces mucus secretion via Cftr mediated pathways, and that in the absence of a functional Cftr channel, PGE_2_ can help release mucus from the goblet cell.

**Fig 4:**
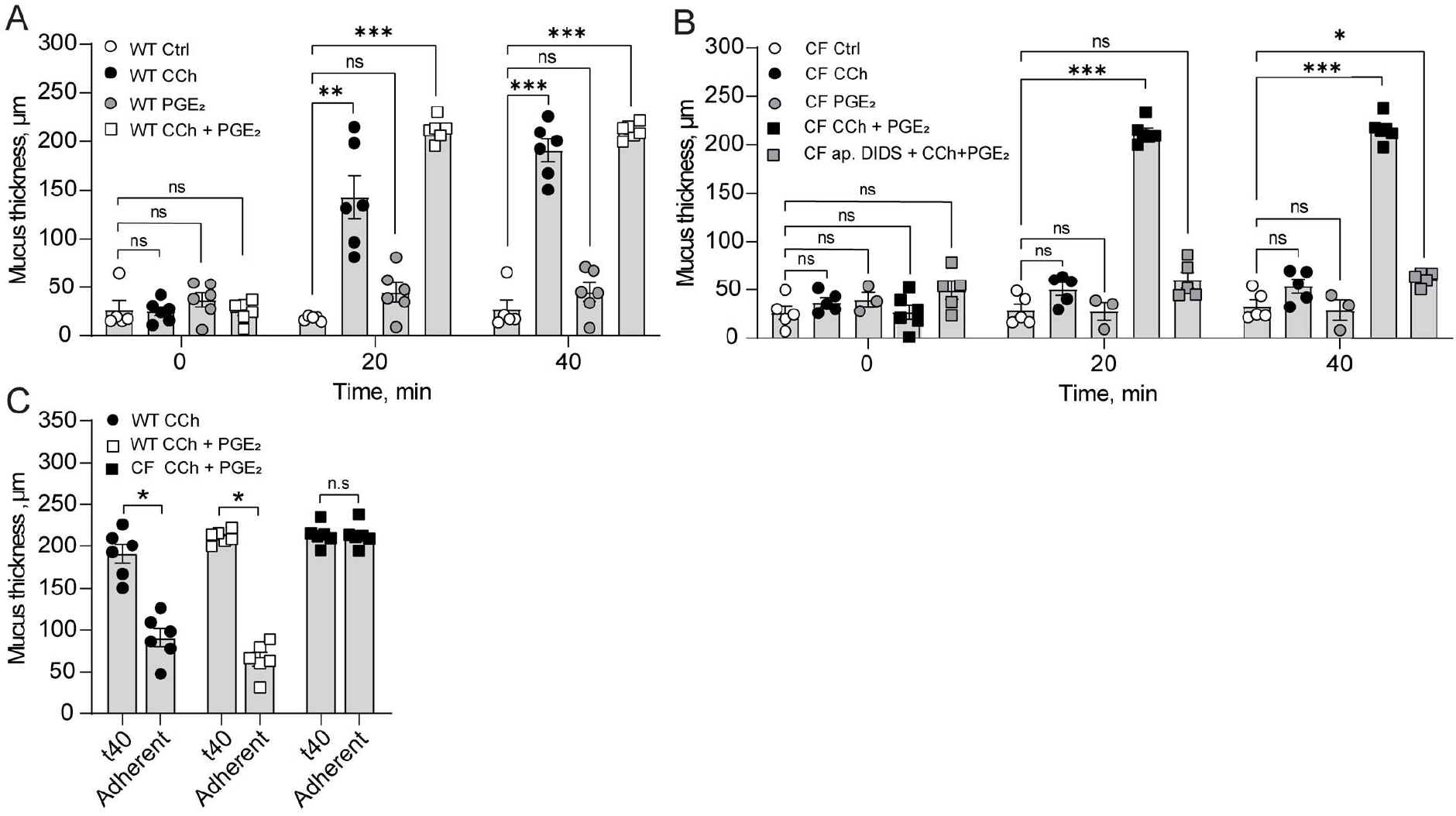
Effect of CCh and PGE_2_ in regulating mucus layer formation in the WT and CF ileum. A) Mucus thickness in WT Ctrl (n=5), WT treated with CCh (10 µM) (n=6), WT treated with PGE_2_ (10 µM) (n=6), and WT treated with CCh and PGE_2_ (10 µM + 10 µM) (n=5). B) Mucus thickness in CF Ctrl (n=5), CF treated with CCh (10 µM) (n=5), CF treated with PGE_2_ (10 µM) (n=3), CF treated with CCh and PGE_2_ (10 µM + 10 µM) (n=6), and CF treated with apical DIDS (200 µM) + CCh (10 µM) + PGE_2_ (10 µM) (n=5). C) Mucus adhesion in WT CCh, WT CCh + PGE_2,_ and CF CCh + PGE_2_. CCh, PGE_2_ and the combination of CCh + PGE_2_ were added directly after the first thickness measurement (t0). DIDS was added 20 min prior to the first thickness measurement. Data are presented as mean ± SEM, ^*^p < 0.05, ^**^p < 0.01, ^***^p < 0.001, n.s = non-significant. Statistical analysis was performed using a two-way ANOVA with Sidak’s post-hoc test (A and B), and a paired Mann Whitney test (C).

## Discussion

In both the small and large intestine, maintaining a functional mucus barrier is essential to maintain homeostasis. In the small intestine, a loose and permeable mucus barrier allows for efficient nutrient absorption and facilitates transport of waste products and microorganism to the colon. In the mid to distal colon, the mucus layer transforms into a physical barrier separating the vast majority of live microbes from accessing the underlying epithelium [15, 17, 29]. How the intestines regulate mucus properties to fit the needs of the respective organs is not fully understood but the ionic milieu in the intestinal lumen is known to play an important role in regulating mucus formation by affecting mucin granule exocytosis, mucin expansion, and mucus adhesion to the epithelium [7, 8, 30]. In this study we discovered that apical anion exchange regulates mucus expansion in the distal colon both during steady state and in response to the secretagogue PGE_2_. Cftr on the other hand was shown to regulate CCh induced *de novo* mucus secretion in both the small intestine and distal colon, and to regulate mucus adhesion in the small intestine.

In the intestines, anions can be transported into the lumen via several transport processes including the anion channels, Cftr and Tem16A, and a number of anion exchangers of which the Cl^-^/HCO_3_^-^ exchangers Slc26a3 and Sc26a6 that are expressed in the colon and ileum, respectively are the most characterized [31-34]. As mentioned previously, studies have shown that Cftr plays an essential role in regulating mucus formation in the small intestine by preventing mucus adhesion to the tissue, and in the context of mucus it is transport of bicarbonate into the intestinal lumen that is particularly important to ensure that the mucus is released and expanded correctly [6, 8, 9, 20, 35]. In the colon, the Ca^2+^ regulated anion channel Tmem16A has been implicated in the regulation of baseline mucus release, but the knowledge regarding the role of additional transporters in regulation of mucus expansion following secretion, and how anion transport is involved in regulating induced secretion is lacking [23]. In the present study we demonstrate that baseline mucus growth in the colon is stimulated by anion exchange and independent of a functional Cftr channel. Since the anion exchange inhibitor DIDS did not affect the mucus content of the tissue, we interpret this to mean that anion exchangers drive expansion of previously secreted mucus rather than stimulating *de novo* mucus secretion. Taken together these findings demonstrate that both Tmem16A and anion exchangers are involved in regulating steady state mucus formation in the colon by stimulating mucus release and mucus expansion, respectively. However, although inhibition of anion exchange by DIDS reduced baseline mucus growth, it was not completely inhibited, pointing towards additional mechanisms being involved in regulating colonic mucus expansion following secretion. Proteolytic processing of the Muc2 has been shown to occur in both the small intestine and colon, which may account for the residual mucus growth observed in the presence of the anion exchange inhibitor DIDS [36, 37]. Due to the fact that the ileum does not exhibit a measurable steady state mucus growth in the *ex vivo* [8, 12] we were not able to evaluate the role of anion exchange in regulation of baseline mucus expansion in the ileum.

In both the small and large intestine, mucus is secreted at a steady state rate but can also be induced by Ca^2+^ mobilizing agents such as acetylcholine and its analogue CCh [7, 38-40]. In both the small and large intestine, exposure to CCh induces a rapid increase in intracellular Ca^2+^ levels which triggers mucin granule exocytosis and activates transcellular anion transport and subsequent fluid secretion [39]. Although the Cftr is not directly regulated by Ca^2+^, activation of Ca^2+^ dependent K^+^ channels will trigger an influx of anions ions that leave the cell via the Cftr [30, 41, 42]. Previous studies have shown that CCh induced mucin granule fusion with the plasma membrane remains intact in CF distal colon tissues, while studies in small intestine organoids from CF mice show that mucus remains within the goblet cells following CCh stimulation [7, 30, 43]. Our results confirm that the same pattern with mucus remaining within the goblet cells following CCh stimulation is observed in intact tissue specimens, which supports a role for Cftr in releasing mucus from the goblet cell *in vivo*.

In addition to testing the role of anion transport in regulation of CCh induced mucus secretion, we evaluated the role of PGE_2_ in regulating mucus layer formation. PGE_2_ is a well-established regulator of mucus and bicarbonate secretion in the stomach where it protects the stomach epithelium from the acidic luminal content [44, 45]. In the intestine, the role of PGE_2_ in regulating mucus secretion appears to be species dependent. In rat colon, PGE_2_ induces *de novo* mucus secretion, and increases mucus output [46]. In contrast, studies in human colonic explants and mouse ileum, have not observed *de novo* mucus secretion in response to PGE_2_, but PGE_2_ has been shown to induce mucus expansion in human colonic primary cultures [9, 47, 48]. In the present study we show that in mouse distal colon, stimulation with PGE_2_ induces an increase in mucus growth rate without signs of *de novo* mucus secretion. We interpret this to mean the PGE_2_ induces expansion of already secreted mucus. In the mouse ileum experiments, PGE_2_ did not stimulate an increase in mucus thickness, which correlates with previous findings demonstrating that PGE_2_ does not induce *de novo* mucus secretion. The observed species differences in the effect of PGE_2_ in regulating *de novo* mucus secretion may be related to species differences in prostaglandin receptor (Ptger) expression. Mouse colonic goblet cells primarily express Ptger4 that signals via cAMP, and to a lower extent Ptger1 that signals via Ca^2+^, and Ptger3 that inhibits cAMP production [49]. A similar pattern is seen in human colonic goblet cells that express low levels of Ptger4, 1 and 3 [50]. Rat colonic goblet cells on the other hand, express all four Ptgers, thus have the possibly to induce stronger Ca^2+^ responses triggering *de novo* mucus secretion [51].

Despite our findings that neither CCh nor PGE_2_ triggered *de novo* mucus secretion in CF ileum and colon when added separately, our results demonstrate that when added simultaneously they trigger *de novo* mucus secretion in both organs. Since the mucus secretory response was inhibited by apical DIDS, or apical Cl^-^ free buffer we interpret this to mean that in situations when *de novo* mucus secretion is triggered by elevated intracellular Ca^2+^ for example by acetylcholine, activation of apical anion exchange can help release mucus from the goblet cells and expand the secreted mucus. However, despite normalizing *de novo* mucus secretion in the CF small intestine, the combination of CCh and PGE_2_ did not reverse mucus adhesion to the tissue. This observation, that activation of parallel secretory pathways, normalizes mucus secretion but not mucus adhesion in the small intestine may explain the more severe mucus pathology in the small intestine as compared to the colon. *In vivo*, Ca^2+^ and cAMP mediated pathways will be active in parallel by substances such as Ach, VIP, histamine, 5-HT, PGE_2_, etc. that all signal via G protein coupled receptors. In the small intestine, this will result in release of adherent mucus, that triggers a vicious circle of mucus secretion, obstructions and inflammation. In the colon on the other hand, Cftr is not involved in regulation of baseline mucus formation, and parallel activation of Ca^2+^ and cAMP mediated pathways can restore Ca^2+^ mediated mucus release from the colonic crypts, protecting them from the microbiota [52].

In conclusion, this study demonstrates that anion transport via the Cftr and apical anion exchange are involved in regulating mucus layer formation in both the ileum and distal colon. Apical anion exchange appears to induce expansion of already secreted mucus, while Cftr drives *de novo* mucus secretion from the ileal and colonic crypts. In the absence of a functional Cftr channel, activation of apical anion exchange can help release mucus from the goblet cell.

## Acknowledgement

This work was supported by the Swedish Research Council (no. 2020-01588 and 2023-02474), The Knut and Alice Wallenberg Foundation (2017.0028), the European Research Council ERC (101100663), Inga Britt and Arne Lundberg foundation (2018-0117), Sahlgren’s University Hospital (ALFGBG-440741, the ALF agreement 236501), Wilhelm and Martina Lundgren’s Foundation, Åke Wiberg’s Foundation, Jeanssson’s foundation, Ollie and Elof Ericsson’s foundation, Apotekare Hedberg’s Foundation, and Riksförbundet för Cystisk fibros.

## Declarations

### Ethics approval

All animal procedures and protocols were performed in accordance with Swedish and EU animal welfare legislations and approved by the Swedish Laboratory Animal Ethical Committee in Gothenburg (Ethical permit ID number: 5.8.18-11053/2019). Animals were anaesthetized by isoflurane inhalation (Isoba® Vet, Schering plough, USA) and euthanized by cervical dislocation and removal of the heart.

### Consent for publication

All authors have reviewed and agree with the manuscript content.

### Conflict of interest

The authors declare that they have no competing interests.

### Author contribution

PLL, AE, MMSG, VLP and JKG performed the experiments and analyzed data. JKG and AE designed the study and interpreted data. JKG wrote the manuscript.

### Data availability

All mentioned data are represented in the main manuscript figures. Other additional data will be made available on reasonable request.

## Notes

### Competing Interest Statement

The authors have declared no competing interest.

